# The Cell Wall Controls Stem Cell Fate in the Arabidopsis Shoot Apical Meristem

**DOI:** 10.1101/2025.05.19.654883

**Authors:** Tasnim Zerin, Paola Ruiz-Duarte, Ann-Kathrin Schürholz, Theresa Schlamp, Yanfei Ma, Carlo Bevilacqua, Nabila El Arbi, Christian Wenzl, Andrej Miotk, Robert Prevedel, Thomas Greb, Jan Lohmann, Sebastian Wolf

**Author notes:** These authors contributed equally to this work (Tasnim Zerin, Paola Ruiz-Duarte, Ann-Kathrin Schürholz).

## Abstract

At the basis of plant developmental plasticity are continuously active pluripotent stem cells, which fuel the life-long post-embryonic formation of new organs. Since plant cells are encased in cell walls and thus immotile, their individual fate-specification program is dependent on their relative position within the organism. It is assumed that the cell wall and its mechanical properties are under surveillance, linking cell wall state to intracellular gene-regulatory networks. However, the role of these cell wall signalling pathways in plant development and the contribution of cell wall properties to cell behaviour and identity are unclear. Here, we show that control of cell wall properties is essential for the perpetuation of stem cell populations and pattering in the shoot apical meristems. The expression of pectin methylesterases (PMEs), which modify homogalacturonan methylesterification and thereby modulate cell wall mechanics, is maintained at low levels in stem cells by the stem cell specifying transcription factor WUS. Low PME expression is required for stemness, auxin patterning and stem cell-specific mechanical properties. Conversely, WUS depletion reduces wall stiffness in the meristem centre, reinforcing its role in maintaining mechanical homeostasis. Together, our findings show that WUS-mediated control of cell wall-modifying enzymes is essential for sustaining stem cell identity and SAM organization. These results demonstrat that the plant cell wall is not only involved in cell differentiation, but also exerts feedback control on developmental transitions, contributing to our understanding how the immediate physical environment is able to guide cell fate decisions in plants.

## Introduction

Plant stem cells pools are localized in regions called meristems, where they are maintained by local signals. In the shoot apical meristem (SAM), which harbours the stem cell niche responsible for most of the above-ground organs, stem cells located in the centre of the meristem dome are specified by the homeobox transcription factor WUSCHEL (WUS), which is expressed in niche cells below the stem cell region but moves to the top layers through plasmodesmata (Fuchs and Lohmann, 2020). Stem cells specifically express the signalling peptide CLAVATA3, which in turn inhibits WUS activity (Brand, 2000; Mayer et al., 1998; Schoof et al., 2000). This negative WUS-CLV3 feedback loop maintains stem cell homeostasis tailored to environmental and developmental cues (Landrein et al., 2018; Pfeiffer et al., 2016; Wenzl and Lohmann, 2023). Stem cell daughter cells divide and are displaced from the meristem centre towards the periphery, where they are recruited to a specific differentiation program such as the formation of a new organ (i.e. leaf or inflorescence). Along their trajectory, cells pass through several transcriptionally distinct domains, demonstrating that here, in contrast to other meristems, cell identity is uncoupled from clonal identity, rendering the SAM an ideal system to study cell fate specification. Interestingly, the various SAM domains differ dramatically in cellular growth dynamics, cell geometry, and mechanics (Kierzkowski et al., 2012; Milani et al., 2011; Peaucelle et al., 2011a; Qi et al., 2017), all of which mainly depend on the composition and remodelling of the plant’s extracellular matrix, the cell wall. In animals, the extracellular matrix forms an integral part of the stem cell niche (Watt and Huck, 2013). Even though the cell wall impinges on almost all aspects of cellular behaviour in plants and is the major constraint for growth, little is known about the contribution of the cell wall to plant cell identity. However, it has been established that enzymatic modification of pectic homogalacturonan, a major component of the cell wall, plays important roles in SAM mechanics and organ production (Peaucelle et al., 2008, 2011a; Qi et al., 2017). Homogalacturonan is synthesized in a methylesterified state and secreted to the wall, where its methyl groups can be removed by the enzyme pectin methylesterase (PME). In the SAM, primordia are associated with pectic epitopes with a low degree of methylesterification and high wall extensibility, whereas the stem cell region shows a high degree of methylesterification and stiffer cell walls (Milani et al., 2011; Peaucelle et al., 2011a). Strikingly, local PME activity can trigger the outgrowth of a new organ, whereas global reduction of PME activity leads to meristems with low cell wall extensibility and devoid of organs, indicating that the state of the pectin plays important roles in meristem maintenance. Here, we show that PME activity is under control of the stem cell specifying transcription factor WUS in the shoot apical meristem, restricting PME expression to the periphery. This insulation of the stem cells to PME activity is essential for maintaining stemness and altering stem cell wall properties has wide-spread effects on cell morphology, tissue patterning and SAM-wide gene expression. Together, our results demonstrate that cell wall properties are essential for stem cell maintenance and that the cell wall state controls cell fate.

## Results

### Stem cells show specific regulation of cell wall properties

Homogalacturonan (HG), a polymer of galacturonic acid subunits is produced and modified inside the cell and secreted to the wall in a highly methylesterified form. In the SAM, pectin de-methylesterification is associated with increased wall softness and precedes organ outgrowth, whereas the stiffer meristem centre, including the stem cells, are characterized by highly methylesterified HG (Milani et al., 2011; Peaucelle et al., 2008, 2011a). Global reduction of methylesterification by expression of the well-characterized PME VANGUARD1 (VGD1) (Jiang et al., 2005; Wolf and Greiner, 2012) leads to reduced SAM size and cell number (Figure 1a-c), but increased cell size (Figure 1d, Supplemental Figure 1), suggesting that spatial control of PME activity is not only important for primordial outgrowth, but also for cell proliferation in the SAM. A query of published cell-type specific expression data (Tian et al., 2014; Yadav et al., 2014, 2009) suggests that PMEs are highly expressed in the SAM periphery but barely expressed in the stem cells (Figure 1e,f,g), consistent with the observed pattern of methylesterification (Peaucelle et al., 2008). In contrast, HG biosynthetic enzymes of the GAUT family of galacturonic acid polymerases (Atmodjo et al., 2011) show similar cumulative expression values across all SAM domains (Figure 1e). The 16 most highly expressed PMEs in the SAM all show low expression in the CLV3 domain, i.e. in stem cells Figure 1f. Interestingly, 10 of the 16 highly expressed PMEs, or 62.5%, appear to be direct targets of the stem cell-specifying transcription factor WUS based on our previous ChIP-seq experiments ((Ma et al., 2019), Figure 1f). RNA-seq analysis revealed that a plurality of SAM-expressed PMEs showed reduced expression after ubiquitous WUS induction (Figure 1f, arrows, (Ma et al., 2019)). In agreement with the known WUS mode of action (Kieffer et al., 2006; Ma et al., 2019), WUS-targeted loci encoding PMEs that showed reduced transcript abundance also generally show a reduction in H3K9/14 acetylation in ChIP-seq data (Ma et al., 2019). In agreement with a largely negative regulation of PMEs by WUS, PME activity was significantly reduced 24 hours after WUS induction in the pUB10:WUS-GR line (Figure 1h). Taken together, these results suggest that negative PME regulation by WUS could explain pectin state in stem cells.

**Figure 1.**
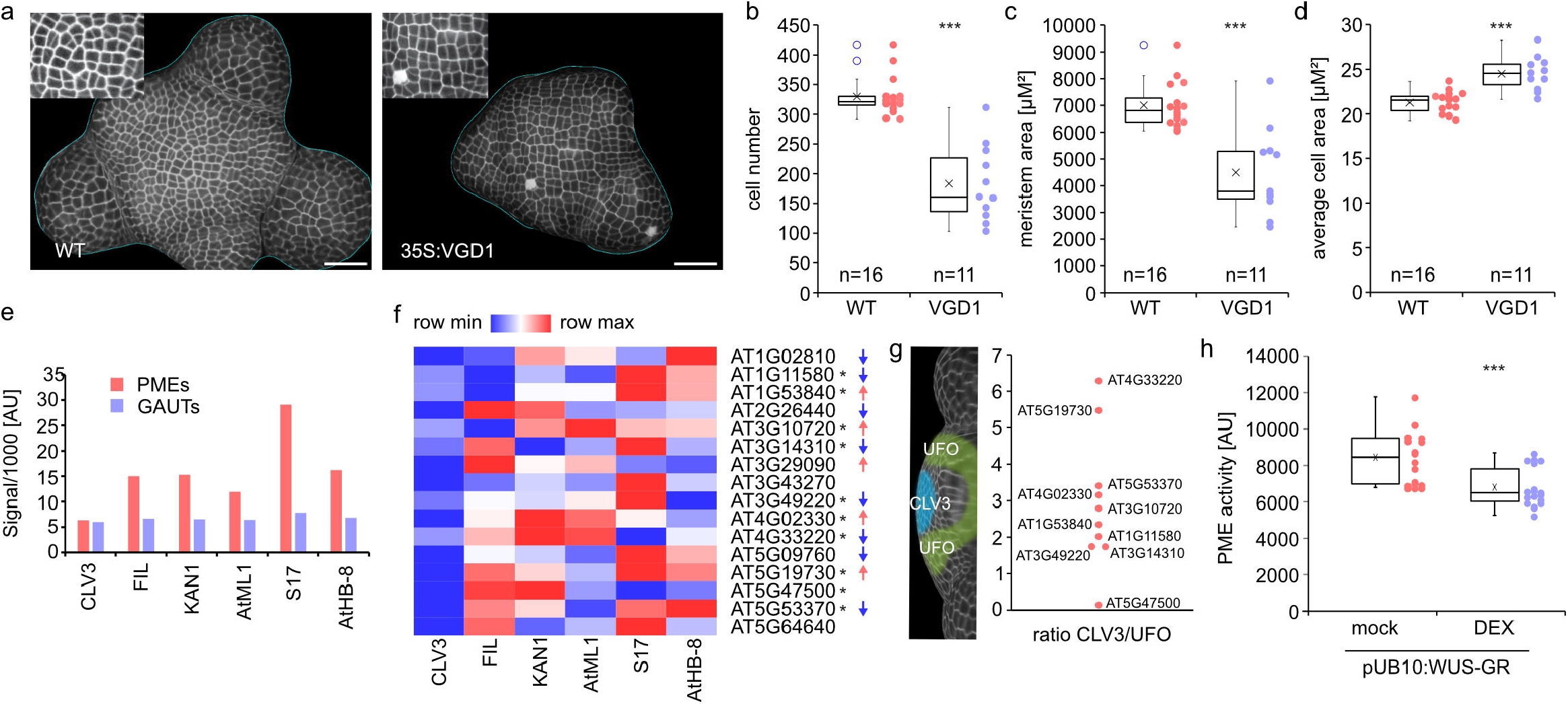
PME expression alters meristem size and cell properties. a) Representative projections of WT (Col-0) and 35S:VGD1 plants showing alterations in shape and size of the meristem as well cells. Inlays= magnification of the central regions. Scale bars = 50 µm. b-d) Quantification of cell number (b), meristem area (c), and average cell area (d) in WT and 35S:VGD1 lines. Graphs depict indicate interquartile range (box), median (bar) and 1.5x interquartile range (whiskers) as well as individual data points. Asterisks indicate statistical significance with P<0.001 according to Student’s t test. e) Summed Microarray data from publishes cell type-specific SAM expression analysis f) heat map showing expression strength of the 16 most highy expressed PMEs in the SAM (based on (Yadav et al., 2014, 2009)) demonstrating low specific expression in the CLV3 domain. Asterisks indicates WUS binding in ChIP seq experiments (Ma et al., 2019), arrows indicate direction of significant differential expression after WUS-GR induction (Ma et al., 2019). g) ratio of expression values in the UFO domain versus the CV3 domain of PME WUS targets in the TRAP-seq experiments described in (Tian et al., 2014). h) PME activity 24 hours after WUS induction in pUB10:WUS-GR seedlings. Data points indicate individual replicates.

### WUS controls PME expression in stem cells and the SAM periphery

To visualize the SAM expression patterns of WUS-targeted PMEs in the SAM, we created a fluorescent reporter line expressing erCitrine under control of the PME PCRF (PMEPCRF, at5g53370) regulatory regions. Fluorescence derived from the pPMEPCRF:erCitrine reporter was absent from the *WUS*-expressing centre of the meristem (Figure 2a). This observation was consistent with the low abundance of PME transcripts in stem cells suggested by cell type-specific transcriptomes (Yadav et al., 2014, 2009). For other PMEs with reduced expression after WUS induction we were able to confirm ChIP-seq results by ChIP-qPCR, demonstrating that presence of WUS leads to a reduction of H3K9/14 acetylation, consistent with a negative regulation of the majority of the SAM-expressed PMEs by WUS (Figure 2b,c, Supplemental Figure 2). This was also consistent with WUS acting primarily as a transcriptional repressor in stem cells and the negative effect of induced ubiquitous activity in plants carrying *pUBQ10:WUS-GR* on the expression of the plurality of SAM-expressed PME genes (Ma et al., 2019). In that context, we were intrigued by a possible regulation of PME5 by WUS, since PME5 is not among the downregulated genes after WUS-GR induction (Figure 1f), but harbours a single canonical WUS G-box binding site (Busch et al., 2010) at position −848 bp upstream of the start codon (Figure 2d), and was implicated in meristem function before (Peaucelle et al., 2011b). In contrast to the majority of PMEs, PME5 did not show WUS-induced changes in H3 acetylation (Figure 2d). A pPME5:erCitrine reporter showed fluorescence broadly across the meristem in a heterogeneous pattern, which was strongly reduced in the central region, consistent with transcriptome data (Figure 2a). In order to directly assess the impact of WUS regulation in the reporter background, we deleted the single WUS binding site using CRISPR/Cas9 in the pPME5:erCitrine line. This deletion, which we confirmed to be present in both the endogenous *PME5* locus as well as in the reporter T-DNA, strongly reduced reporter expression in the periphery of the SAM (Figure 2e), suggesting WUS positively regulates expression of this PME. This was corroborated by reduced expression in lines expressing a construct in which the WUS G-box was mutated (Figure 2f), suggesting that reduced expression in the CRISPR deletion line was not primarily caused by adjacent, WUS-independent motifs. Conversely, WUS induction in the WUS-GR line enhanced pPME5:erCitrine-derived fluorescence, corroborating positive regulation by WUS (Figure 2g, Supplemental Figure 3). Notably, reporter expression was still low in the centre of the meristem and increased only at the periphery. Inspection of WUS-GR genomic data revealed a decrease of the repressive H3K27 trimethylation epigenetic mark at the *PME5* locus, consistent with a positive effect of WUS on *PME5* expression(Figure 1e,f)(Ma et al., 2019). In agreement, ChIP-qPCR experiments using an anti-H3K27me3 antibody showed strongly increased abundance of *PME5* DNA in the CRISPR deletion line compared to the background line without the deletion (Figure 2fh) demonstrating that WUS enhances PME5 expression by reducing histone methylation at its locus. In summary, PME expression is low in stem cells and WUS contributes to PME regulation through multiple mechanisms involving various epigenetic modifications.

**Figure 2.**
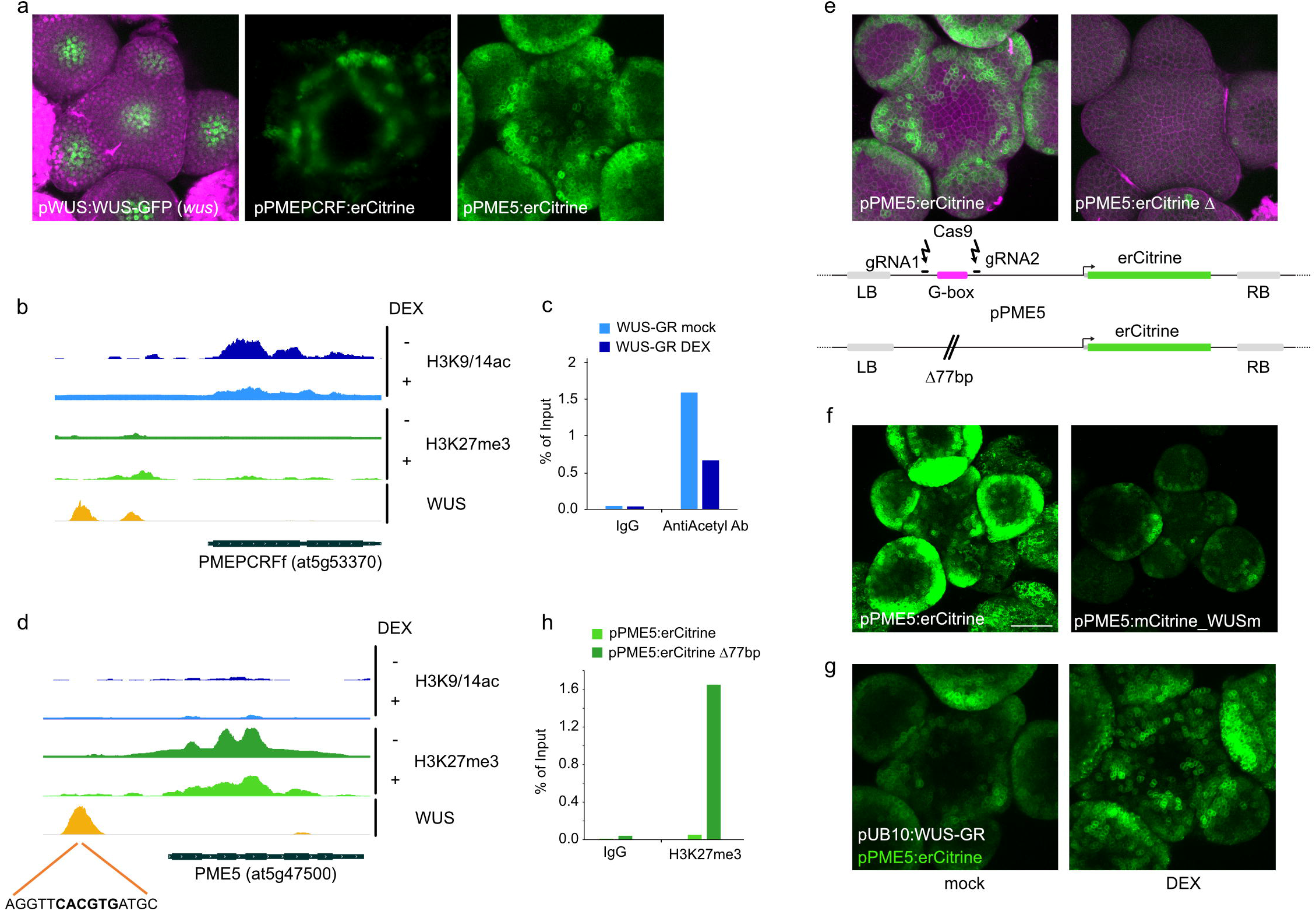
WUS restricts PME expression in the stem cell region of the SAM. A) representative maximum projections of z-stack images of the indicated lines. b) CHiP-seq tracks (Ma et al., 2019) of the PMEPCRF genomic region before and after WUS-GR induction for Anti-H3ac (blue), Anti-H3K27me3 (green) and anti-WUS. c) Representative ChIP-qPCR conformation of reduced H3 acetylation at the *PMEPCRF* locus, see Supplemental Figure 2 for replicates. d) same as in b) but for PME5, see Supplemental Figure 2 for replicates. e) Effect of CRISPR-Cas9-mediated deletion of the WUS binding motif region in the pPME5:erCitrine line. f) Effect of mutation of the G-box in the pPME5:erCitrine construct. g) WUS induction in the WUS-GR line enhances the pPME5:erCitrine fluorescence. h) ChIP-qPCR showing that deletion of the WUS binding site in the pPME5:erCitrine line targeting both reporter and endogenous PME locus, increases H3K27me3. For replicates see Supplemental Figure 2.

### Control of cell wall properties is required for stem cell function

To determine the significance of the low PME expression in stem cells, we tested the effect of counteracting the endogenous regulation by inducing PME activity specifically in CLV3-positive cells. To this end, we made use of our inducible cell type-specific trans-activation system (Schürholz et al., 2018). We used a driver line expressing the synthetic transcription factor LhG4 fused to the ligand-binding domain of the rat glucocorticoid receptor under control of the *CLV3* promoter. In the absence of the inducer Dex, the chimeric TF was retained in the cytosol and only translocated to the nucleus after induction. In the nucleus, LhG4 drove expression of genes under control of a cognate synthetic promoter (pOp6) (Craft et al., 2005; Moore et al., 1998), in this case a ER-directed mTurquoise2 fluorescent reporter ((Schürholz et al., 2018), Figure 3a). We crossed this driver line to an effector line carrying the previously described PME *VGD1* under control of the pOp6 promoter and analysed doubly homozygous material in comparison to the original driver line (Figure 3a). To confirm that PME expression in stem cells altered physical properties of the cell walls in that region, we performed Brillouin microscopy coupled to a confocal setup. Brillouin microscopy is capable of measuring the mechanical, i.e. visco-elastic properties of biological samples in a non-contact, label-free and high-resolution fashion using light scattering (Prevedel et al., 2019). We quantified Brillouin shift in three regions of the meristem: periphery spanning approximately three cell layers starting with L1 at a 45° angle from the centre line (ROI1), an apical region at the centre of the meristem corresponding to the stem cell region (ROI3), as well as the subtending region, corresponding approximately to the organizing centre (ROI2) (Figure 3b). Brillouin shift measurements indicated that the centre of the meristem was the stiffest of the three regions. This result was in line with AFM measurements (Milani et al., 2011; Peaucelle et al., 2011a) validating our approach. In line with expectations, *VGD1* expression for 48 hours led to an increase in the Brillouin shift in the stem cell region, indicative of reduced stiffness, whereas the other regions were largely unaffected (Figure 3b). In agreement with reduced stiffness and with what was observed with constitutive *VGD1* expression (Figure 1d), *VGD1*-expressing cells in the meristem centre showed a cell area distribution in the L1 layer that was shifted towards larger cells (Supplemental Figure 4). Thus, induction of PME activity in stem cells resulted in altered cell wall properties and mechanics in the central region of the meristem, which translated into modified cell size.

**Figure 3.**
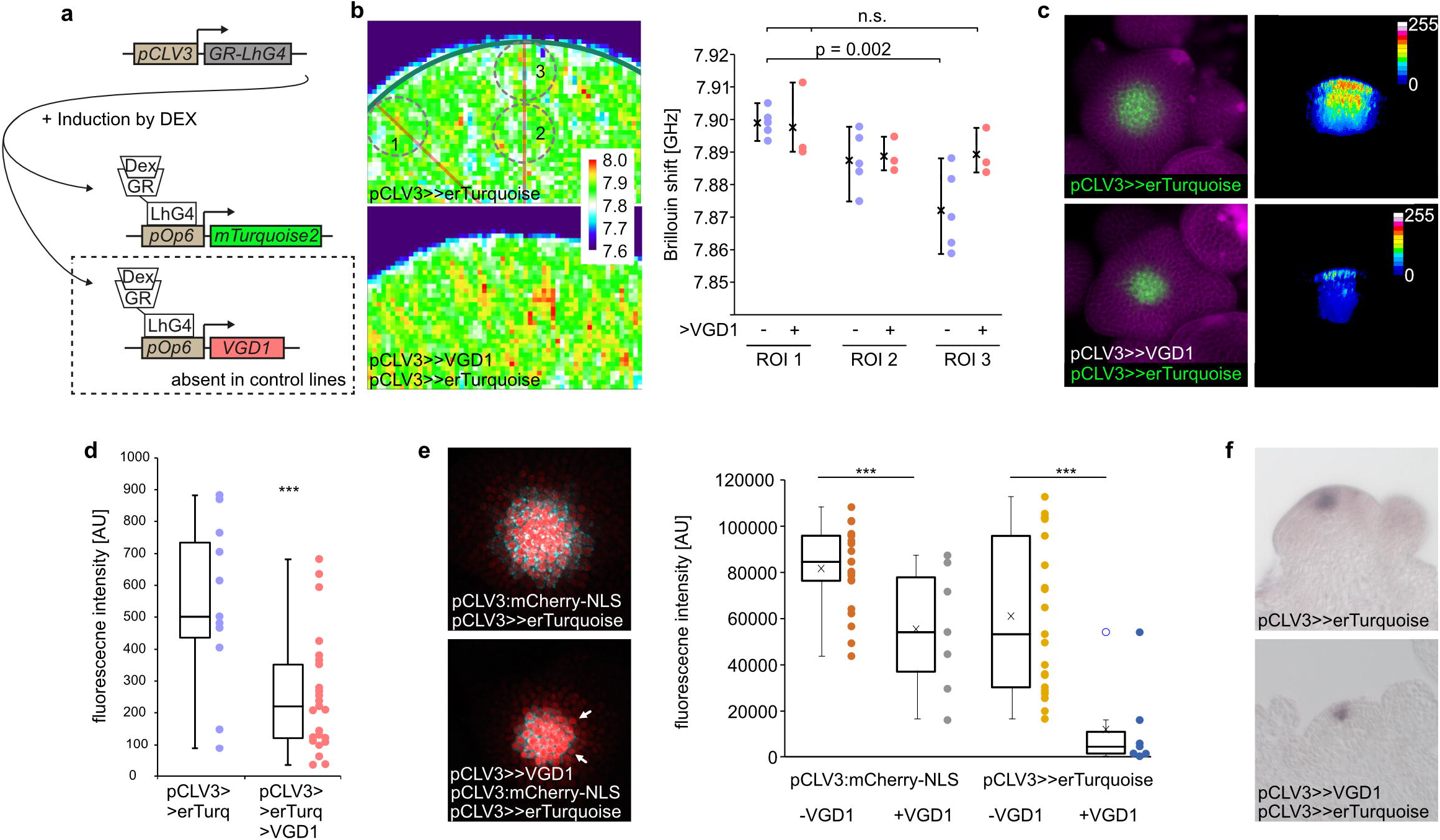
PME activity in stem cells is detrimental to stemness. a) Description of the inducible cell type-specific trans-activation system (Schürholz et al., 2018). The synthetic GR-LhG4 transcription factor is expressed under control of the pCLV3 promotor. Upon DEX-induction GR-LhG4 translocates to the nucleus where it binds to the cognate pOp6 promoter driving expression of ER-directed mTurquoise2 in the control line and both ER-directed mTurquoise2 and a PME (VGD1 or PME5) in the experimental line. Note that the pOp6:PME constructs where transformed into the control line background. b) Brillouin microscopy after VGD1 trans-activation shows increased Brillouin shift indicative of increased elasticity in the central region of the meristem. c) trans-activation (indicated by “>>“) of VGD1 reduces intensity and domain size of pCLV3>>erTurquoise-derived fluorescence. d) Quantification of pCLV3>>erTurquoise-derived fluorescence intensity as in b). e) A pCLV3:mCherry-NLS marker also shows reduced fluorescence and domain size after trans-activation of VGD1. f) RNA-in-situ hybridization of meristems as in b) with a probe against CLV3.

To study the effect of PME induction on stem cell identity, we monitored fluorescence from the pOp6:driven erTurquoise reporter that marks the *CLV3* expression domain (Figure 3c). Induction of VGD1 expression in the stem cells for 72 hours led to a reduction of both intensity and domain size of pCLV3->erTurquoise, suggesting a reduction of stemness in the apical cells of the meristem (Figure c,d, Supplemental Figure 5a). This was corroborated by analysis of a pCLV3:mCherry-NLS reporter crossed into the same lines (Figure 3e, Supplemental Figure 4b). Moreover, mRNA in situ hybridization experiments demonstrated that endogenous *CLV3* transcripts also showed reduced abundance after PME induction (Figure 3f). These effects were not limited to expression of *VGD1*, as a similar apparent reduction in stemness was observed after stem cell-specific expression of *PME5* (Supplemental Figure 5c). Importantly, these effects are unlikely to be due to reduced WUS mobility as we observed normal WUS-GFP fluorescence in L1 and L2 layers of a pWUS-WUS-GFP (*wus*) rescue line (Daum et al., 2014) 72 hours after VGD induction, indicating that PME expression did not interfere with WUS movement. In fact, WUS-GFP fluorescence was present in stem cells that did not show any pCLV3>>erTurqouise signal (Supplemental Figure 6). Together, these data suggested that cell wall properties in stem cells need to be tightly controlled to maintain stemness. Conversely, cell wall properties exert feedback control on cell fate in the shoot apical meristem.

### Stem cell wall properties affect meristem patterning and cause meristem-wide gene transcriptional rearrangements

To assess whether this apparent reduction in stemness caused by stem cell-specific PME expression also caused non-cell autonomous effects, we analysed the auxin response reporter DR5v2:erVenus (Ma et al., 2019) and observed an increase in the size of the auxin response maxima as well as a reduction in size of the auxin response minimum in the centre of the meristem (Figure 4a, Supplemental Figure 7). To study meristem-wide effects of cell wall changes in stem cells, we performed RNA-seq experiments on dissected meristems 72 hours after stem cell-specific PME expression. Of 715 differentially expressed genes (Padj <0.05), 140 were increased and 575 were reduced in expression after *VGD1* induction (Supplemental Dataset 1). Notably, GO enrichment analysis showed that genes associated with leaf and shoot development were overrepresented among the upregulated genes, whereas downregulated genes were enriched in photosynthesis-associated genes. Overall, transcription factors were overrepresented among the differentially expressed genes. As *WUS* and *CLV3* are represented with only very few reads, we reasoned that stem cells were too diluted in the samples to account for significant changes in the meristem transcriptomes. Thus, these results suggested that change in cell wall properties in stem cells can have pronounced meristem-wide effects. This was exemplified by a reporter we created using the promotor of AGAMOUS-LIKE 22/SHORT VEGETATIVE PHASE (AGL22/SVP), a gene that is involved in floral transition (Hartmann et al., 2000) and was among the significantly upregulated genes after *VGD1* induction. Under control conditions, signal derived from pSVP:3xGFP was weak in the shoot apical meristem and restricted to the meristem centre. In contrast, after PME induction in the CLV3 domain, reporter expression could be observed throughout the meristem, corroborating the RNA-seq results and indicating a non-cell autonomous effect of cell wall changes in the stem cells.

**Figure 4.**
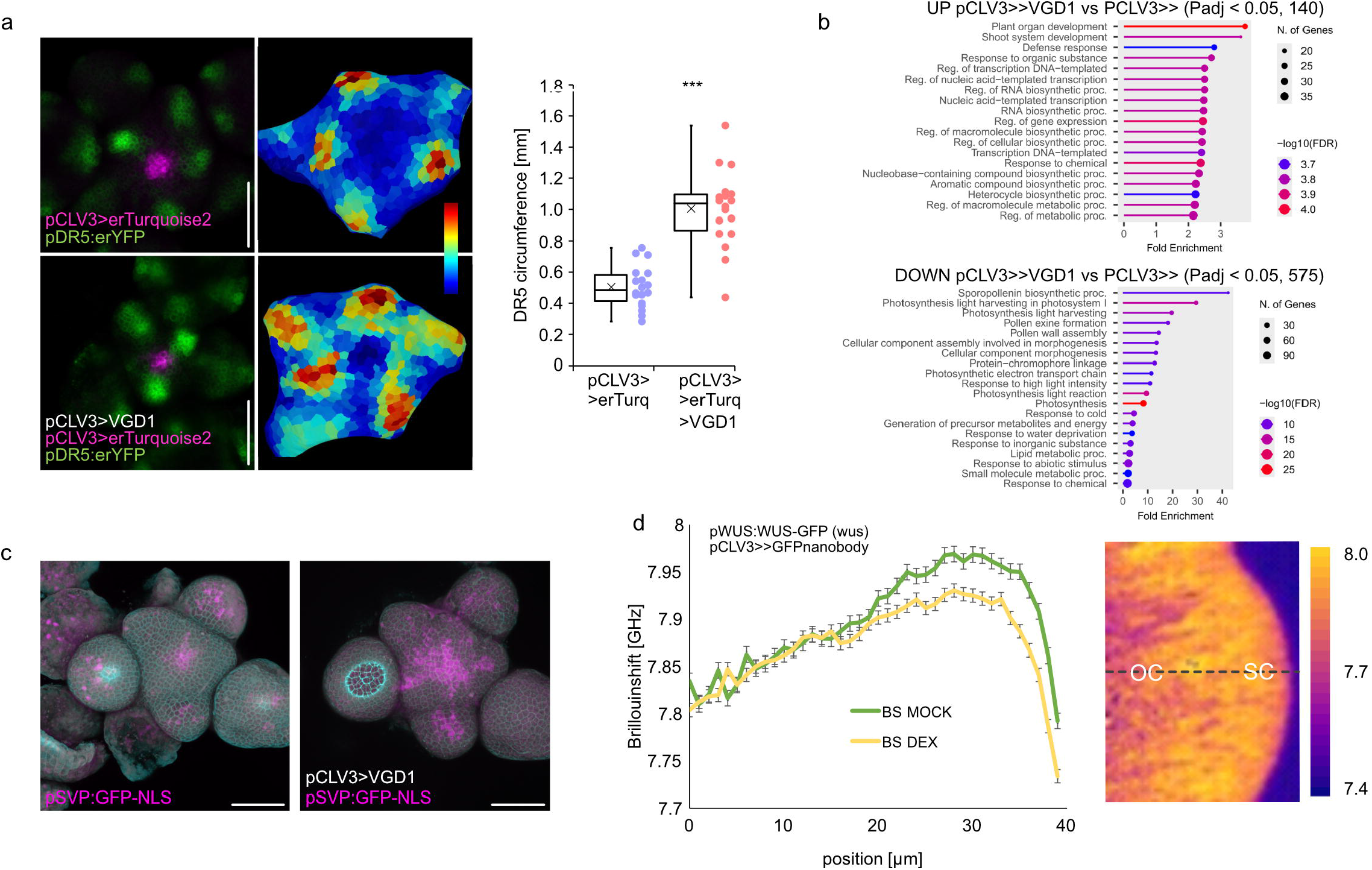
Cell wall properties in the stem cells exert meristem-wide effects. a) PME induction in stem cells leads to an increase in the size of auxin response domains as indicated by the DR5:erVenus reporter. b) GO enrichment analysis of differentially expressed genes according to RNA-seq analysis of dissected meristems 48h after pCLV3>>VGD induction. c) Analysis of pSVP:GFP-NLS expression after pCLV3>>VGD1 induction. d) Brillouin microscopy of WUS rescue line meristems (pWUS:WUS-GFP (wus)(Daum et al., 2014)) with or without DEX induction of WUS degradation with the degradFP system (Caussinus et al., 2012). Results are derived from three independent experiments, each with at least 6 meristems for each condition.

### WUS controls stem cell mechanical properties

Our results showed that cell wall genes, exemplified here by PMEs, are under control of the stem cell-specifying transcription factor WUS. If this hypothesis were correct, we expected cell wall properties to change in the absence of WUS. To test this hypothesis, we made use of the degradFP system (Caussinus et al., 2012) in a WUS-GFP rescue line (pWUS-WUS-GFP (wus) (Daum et al., 2014)). DegradFP is based on nanobodies directed against fluorescent proteins and targeting those for degradation. Placing the degradFP anti-GFP nanobody (NSlmb-vhhGFP4) under control of the pCLV3> driver in the WUS-GFP rescue line thus allows inducible WUS degradation specifically in stem cells. We performed Brillouin microscopy on meristems 72 hours after WUS degradation compared to mock-treated meristems. In agreement with our hypothesis, WUS degradation resulted in a decreased Brillouin shift of the meristem centre, consistent with a decreased stiffness in stem cell walls and thus a reduction in the gradient of mechanical properties between stem cells and the organizing centre.

In summary, our results demonstrate that i) PMEs and cell wall properties are under strict control of the stem cell-specifying transcription factor WUS: ii) tight regulation of cell wall properties is required to maintain SAM patterning; iii) interference with stem cell wall properties has non-cell autonomous effects in the SAM; and iv) maintenance of cell wall properties is required for stem cell fate.

## Discussion

The cell wall is a defining feature of plant morphogenesis and controls cellular growth in a balance with turgor pressure. In addition, it is now widely accepted that cell wall properties are under surveillance to relay information on the biochemical and mechanical state of the cell wall to the cell interior (Chaudhary et al., 2025, 2020; Duan et al., 2020; Dünser et al., 2019; Feng et al., 2018; Lin et al., 2021; Liu et al., 2024; Shih et al., 2014; Wolf, 2022; Wolf et al., 2014, 2012). Plant cell wall properties are an integral aspect of cell differentiation and cell wall biosynthesis and modification enzymes are under tight control of developmental gene-regulatory networks. However, despite the relevance of the cell wall for plant development, few studies have addressed how the cell wall impinges on cell fate (Chaudhary et al., 2021; Wolf et al., 2014). This in sharp contrast to, for example, solid evidence that the extracellular matrix is involved in cell fate acquisition in animals (Engler et al., 2006; Saha et al., 2008). In addition, a seminal study in Fucus brown algae has shown that cell wall components can specify cell fate in this system (Berger et al., 1994). The cell wall of brown algae strongly differs from that of plants (Popper et al., 2011) but similar requirements and challenges for the relay of information to the cell interior are conceivable. Our results demonstrate that plant stem cells require a specific cell wall constitution to retain stemness, consistent with conceptually similar, albeit molecularly unrelated findings in other systems. An open question is whether signalling from the stem cell wall is biochemical or mechanical in nature, or, as suggested by examples from the animal literature, mediated by a feedback loop involving both (Bailles et al., 2019; Hannezo and Heisenberg, 2019; Heisenberg and Bellaïche, 2013; Lecuit and Yap, 2015). In any case, conversion into biochemical signalling must occur at some point during signal transduction, and future research should focus on identifying the cell wall signalling components mediating cell identity maintenance.

Many studies have contributed to establish the importance of WUS in specifying stem cell fate, and a growing number of targets provide insight into the molecular mechanisms of its role in maintaining pluripotency and preventing differentiation. Here, we provide evidence that one of these roles is the control of cell wall modifying enzymes, ensuring the maintenance of cell wall properties in stem cells, which in turn affects SAM mechanics. WUS is known to recruit histone deacetylases through TOPLESS, which in turns leads to a decrease of histone H3 acetylation and thereby mediates repression of WUS target genes (Galvan-Ampudia et al., 2020; Kieffer et al., 2006; Long et al., 2006; Ma et al., 2019). Consistent with this canonical mode of action, all downregulated PME targets of WUS also show de-acetylation after WUS induction (Ma et al., 2019). Our results also suggest that WUS can non-cell autonomously promote the expression of *PME5* through affecting another epigenetic modification, the repressive H3K27me3 mark. In fact, a minor, but considerable fraction of WUS target loci (634 out 6740) showed changes is H3K273me levels in previous ChIP-seq experiments (Ma et al., 2019). Epigenetic modifications have been suggested to serve as a long-term memory for cells even after cells have left morphogenetic fields or other cell fate-determining positional signals (Birnbaum and Roudier, 2017). In particular, H3K27 trimethylation can be maintained through replication (Alabert et al., 2015; Jiang and Berger, 2017), and is thus a plausible mechanism for the non-cell autonomous effects on PME expression in the periphery of the SAM.

## Acknowledgments

This work was supported by the German Research Foundation (DFG) through research grant WO 1660/11-1 to Swand the research grants LO 1450/7-2 and GR 2104/6-2 as part of the research unit FOR2581 “Quantitative Morphodynamics of Plants”. Work in the TG lab was furthermore supported by the ERC consolidator grant PLANTSTEMS and the Innovation Campus Heidelberg Mannheim Health & Life Sciences. The authors thank Guido Grossmann and Phillip Denninger for providing published material.

---

**Supplemental Figure 1**. Cell size distribution frequencies of Col-0 and 35S:VGD1 meristem cells. Data are from 7 meristems each, with 2281 (Col-0) and 1464 (35S:VGD1) individual cells, respectively.

**Supplemental Figure 2**. ChIP-qPCR analysis of PME3, PME34, and PMEPCRF using Anti-Acetyl-Histon-H3 antibody. The relative abundance of target DNA is expressed as a percentage of total input chromatin DNA.

**Supplemental Figure 3**. Time course of mock and DEX-induced pUB10:mCherry-GR-WUS plants expressing the pPME5:erCitrine marker

**Supplemental Figure 4**. Size distribution frequencies of central region cells pCLV3>> driver lines and pCLV3>>VGD1 lines. Data are from 6-7 meristems each, with 583 (pCLV3>>) and 762 (pCLV>>VGD1) individual cells, respectively.

**Supplemental Figure 5**. Effects of PME expression in stem cells. a) Representative meristem projections after MorphographX segmentation and projection of the pCLV3>>erTurquoise signal. Meristems were images 72 hours after DEX induction. b). a) Representative meristem projections after MorphographX segmentation and projection of the pCLV3:mCherry-NLS signal. Meristems were images 72 hours after DEX induction. c) Effect of pCLV3>>PME5 expression on mTurquoise signal and domain size.

**Supplemental Figure 6**. Cross sections of pWUS:WUS-GFP signal in the indicated bakgrounds. Lines were generated by crossing the WUS rescueline described in (Daum et al., 2014) with the pCLV3>>erTurquoise and the pCLV3>>erTurquoise >>VGD1 lines, respectively. Plants were homozygous for all loci.

**Supplemental Figure 7**. Effects of PME expression in stem cells. a) Representative meristem projections after MorphographX segmentation and projection of the pDR5:erCitirne signal. Meristems were images 72 hours after DEX induction.

**Supplemental Dataset 1**. Differentially expressed genes in pCLV3>>VGD1 meristems 48 hours after induction compared to pCLV3>> control lines.

**Supplemental Table 1**. Oligos used in this study.

**Supplemental Table 2**. GreenGate cloning modules used in this study.

## Material and Methods

### Plant material and growth conditions

Plant material used in this study is described in Supplementary Table 1. For SAM imaging, plants were grown in long day conditions (16h light, 8h dark, 22°C, 60-70 % humidity, 100-125 μmol·m-2·s-1 light intensity) for 7 days on half-strength MS plates with 0.9% agar and 1 sucrose before being transferred to soil. Imaging was performed on inflorescence meristems of 15 cm tall stems. Induction of GR-LhG4-mediated expression was performed by spraying shoot apices with 30 µM dexamethasone solution. Inflorescence meristems were dissected by cutting the stem with and primordia were removed by forceps or a canula up to flower stage 3-4. Shoot apical meristems were counter stained, if appropriate, with PI (200 μg/mL) dissolved in water for 5 min and mounted in a small petri dish with 3 % agarose and covered with water.

### Generation of transgenic lines

CRISPR-Cas9-mediated deletion of the WUS binding motif region in the PME5 promoter was performed according to (Wang et al., 2015) using the pHEE401E vector. gRNA target sites were selected based on CHOPCHOP v2 (https://chopchop.cbu.uib.no/) (Labun et al., 2016), primers are listed in Supplemental Table 2. All other constructs used for generating transgenic lines in this study were obtained through GreenGate cloning as previously described (Lampropoulos et al., 2013). Modules and oligo sequences can be found in Supplemental Tables 2 and 3. Previously published modules (Denninger et al., 2019; Lampropoulos et al., 2013) are indicated. The DEX inducible WUS-degradFP line was generated by bringing the anti-GFP nanobody coding sequence (NSlmb-vhhGFP4)(Caussinus et al., 2012) under control of the LhG4-GR transcription factor by the pOp6 promoter.

### Imaging and Image analysis

Shoot apices were imaged with either an upright Nikon A1 Confocal equipped with with a CFI Apo LWD 25x water immersion objective or an upright Leica SP8 with x40 long working distance water objectives. mTurquoise2 fluorescent was excited with a 405 nm laser line and collected between 425 and 475 nm. GFP and mCitrine fluorescence were excited with 488 nm and collected between 500 and 550 nm. PI and mCherry fluorescence were excited with a 561 nm laser line and collected between 570 and 620 nm. All non-quantitative images were processed using ImageJ. MorphoGraphX was used to quantify cell surface area and cell numbers in the epidermis of the SAM (Barbier de Reuille et al., 2015). The original confocal z-stack images were processed using ImageJ Fiji (https://imagej.net/software/fiji/) to generate .tiff files, which were then analyzed in MorphoGraphX 2.0 (https://morphographx.org/) software. The global shape of the sample was extracted as a 2.5D curved mesh which was then segmented into cells by projecting the cell membrane signals onto the shape. This segmented mesh was used to quantify cell size using the process “Mesh/Heat Map/Heat Map”. Gene expression was quantified by calculating the intensity of a marker for each segmented cell.

Brillouin Microscopy was performed as described in (Bevilacqua et al., 2023). RNA in situ hybridization of CLV3 transcripts was performed as described (Schuster et al., 2014).

### Transcriptome analysis

Shoot apices of 15 cm tall stems were DEX-treated in staggered fashion and dissected and frozen in liquid nitrogen after 48 hours. RNA was extracted using GeneMATRIX Universal RNA/ miRNA Purification Kit (Cat. No. E3599 Distributor Roboklon GmbH). Reads were aligned with RNA STAR (Dobin et al., 2013)using default setting. feautureCounts (Liao et al., 2014) was used to tally read numbers from the BAM files using TAIR10.42.gtf as gene annotation file. DESeq2 (Love et al., 2014) was used to assess differential gene expression was assessed by). Go enrichment was determined using ShinyGO (Ge et al., 2020).

### ChIP-qPCR

For the ChIP experiment with inflorescence meristems, 80 meristems were dissected and collected in 1X PBS buffer. After decanting the PBS buffer, 1% formaldehyde was added to the sample and vacuum infiltration was carried out for 20 minutes. Tissue crosslinking was stopped by decanting formaldehyde and vacuum infiltrating the tissue with 0.125M glycine for 5 minutes. Meristems were then rinsed, dried, and frozen. The frozen samples were grounded and gradually mixed with Nuclei Extraction Buffer (NEB). The sample slurry with NEB was filtered twice through Miracloth (Merck Millipore) and liquid sample was extracted as much as possible. This liquid sample was centrifuged at 4000g for 10 minutes. After discarding the supernatant, the pellet was resuspended with NEB in the presence of 0.1M PMSF. And centrifuged at 4°C for 5 minutes at 4000g. The supernatant was removed and the pellet was resuspended in Nuclei Lysis Buffer (NLB). The sample was sheared into fragments of 200– 400 bp using a sonicator (Covaris E220evolution Focused-ultrasonicator). The sonicated sample was centrifuged at 20000g for 5 minutes at 4°C. The supernatant was then transferred to new tubes and at this step, 2-5% of the sonicated sample was saved as input. Each sample was diluted with ChIP dilution buffer and the samples were then incubated with IgG (Normal Rabbit IgG, Sigma-Aldrich) and appropriate antibody at 4°C overnight (O/N) with rotation. Protein G Dynabeads (Invitrogen) were washed three times with ChIP dilution buffer (without protease inhibitors). The beads were added to the O/N incubated samples and immunoprecipitants were collected by rotating the samples at 4°C for 3 hours. Using a magnetic rack, the beads were reclaimed afterwards and washed with low salt buffer, high salt buffer, LiCl wash buffer and TE buffer consecutively. After the last wash, the IP was eluted by suspending the beads in elution buffer. The input DNA control and all the IP samples were then subjected to reverse crosslinking by adding 5M NaCl and O/N incubation at 65°C with shaking. The samples were incubated at 45°C for 2 hours in the presence of proteinase K (20 mg/mL) before DNA recovery. The DNA samples were purified using MinElute PCR Purification Kit (QIAGEN) according to the manufacturer's protocol. QPCR was performed as described in (El Arbi et al., 2024).

## Data availability

The datasets produced in this study are available at GEO as GSE296557.

