## Supplemental Figures 1-7 for "The Cell Wall Controls Stem Cell Fate in the Arabidopsis Shoot Apical Meristem"

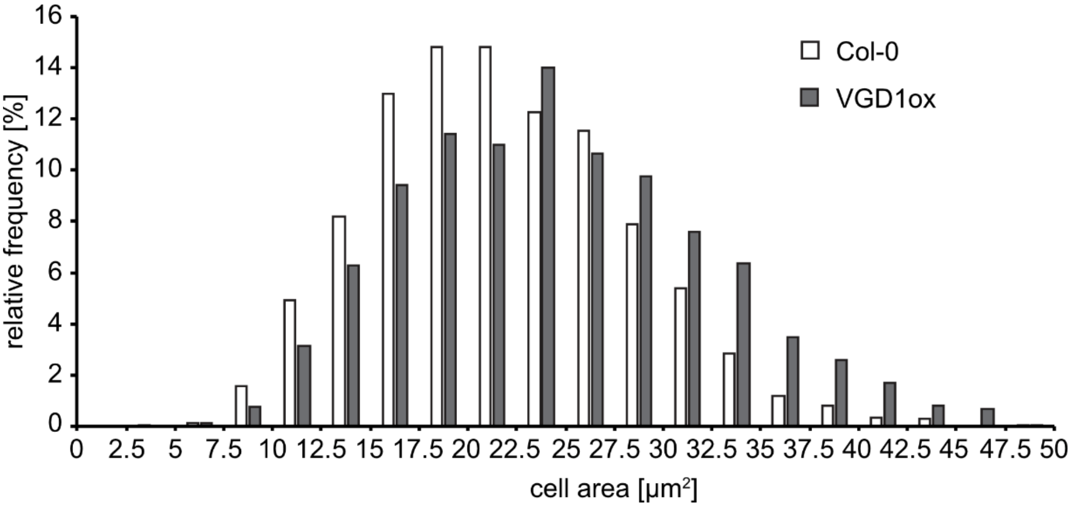


**Supplemental Figure 1.** Cell size distribution frequencies of Col-0 and 35S:VGD1 meristem cells. Data are from 7 meristems each, with 2281 (Col-0) and 1464 (35S:VGD1) individual cells, respectively.

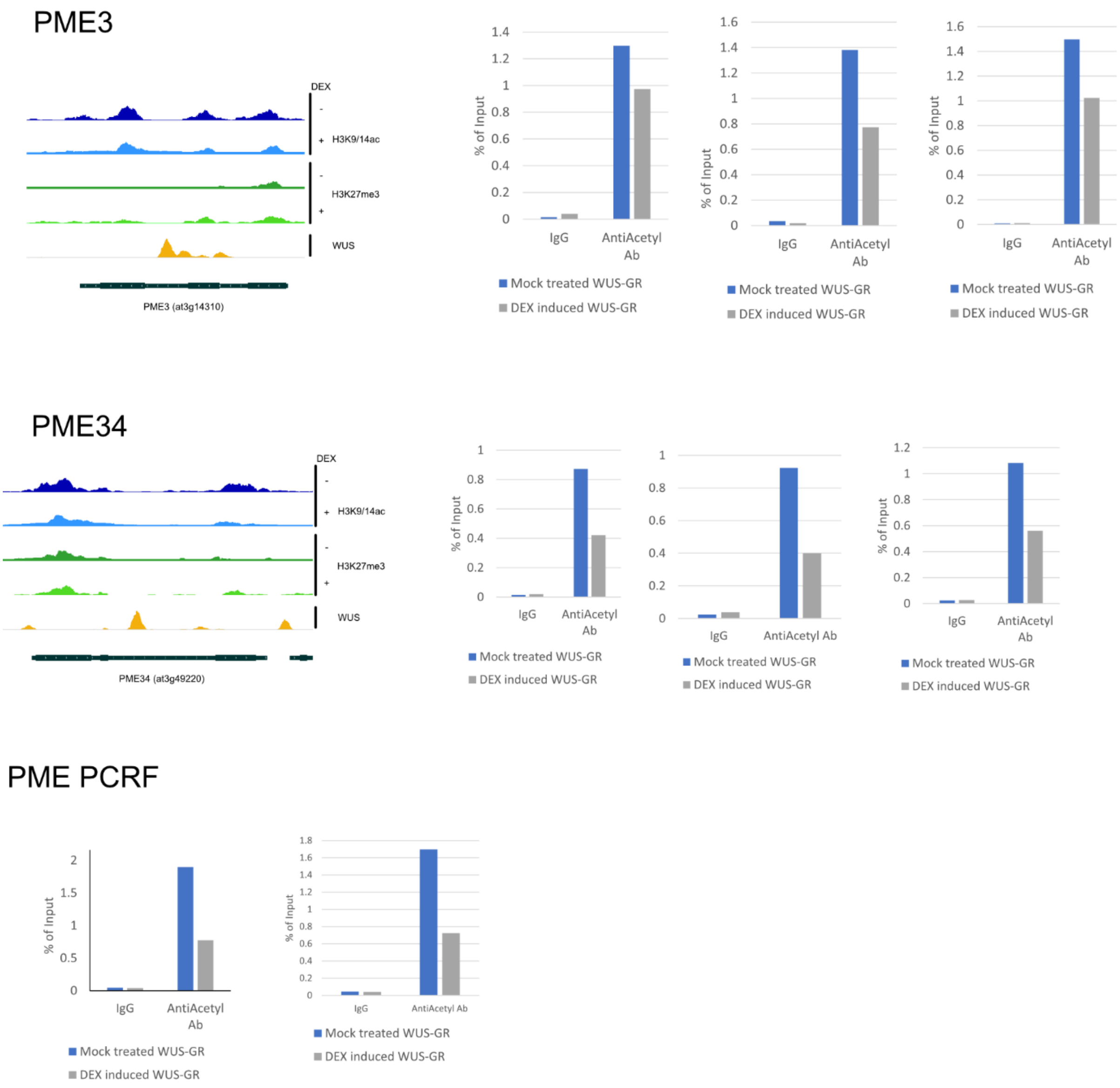


**Supplemental Figure 2.** ChIP-PCR analysis of PME3, PME34, and PMEPCRF using Anti-Acetyl-Histon-H3 antibody. The relative abundance of target DNA is expressed as a percentage of total input chromatin DNA.

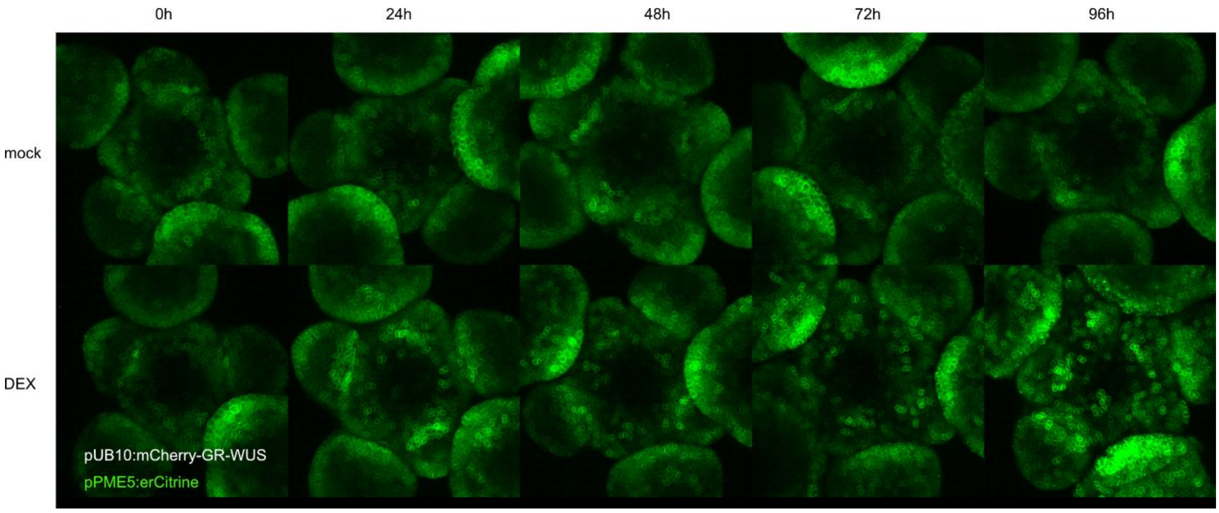


**Supplemental Figure 3.** Time course of mock and DEX-induced pUB10:mCherry-GR-WUS plants expressing the pPME5:erCitrine marker

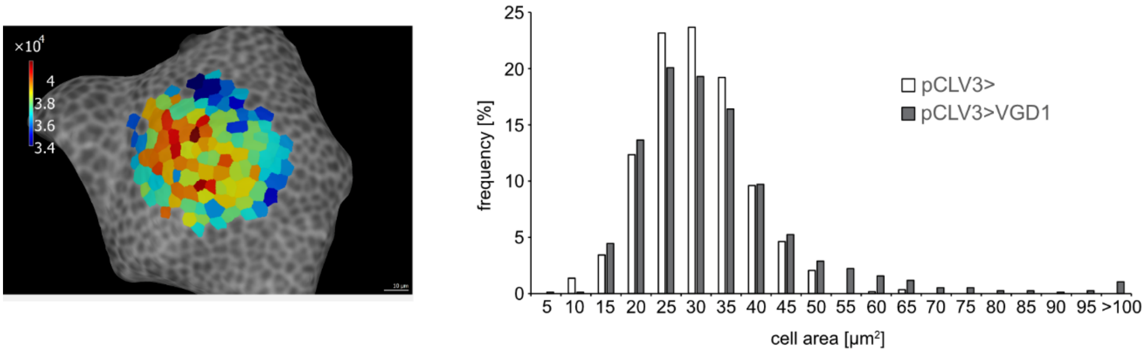


**Supplemental Figure 4.** Size distribution frequencies of central region cells pCLV3>> driver lines and pCLV3>>VGD1 lines. Data are from 6-7 meristems each, with 583 (pCLV3>>) and 762 (pCLV>>VGD1) individual cells, respectively.

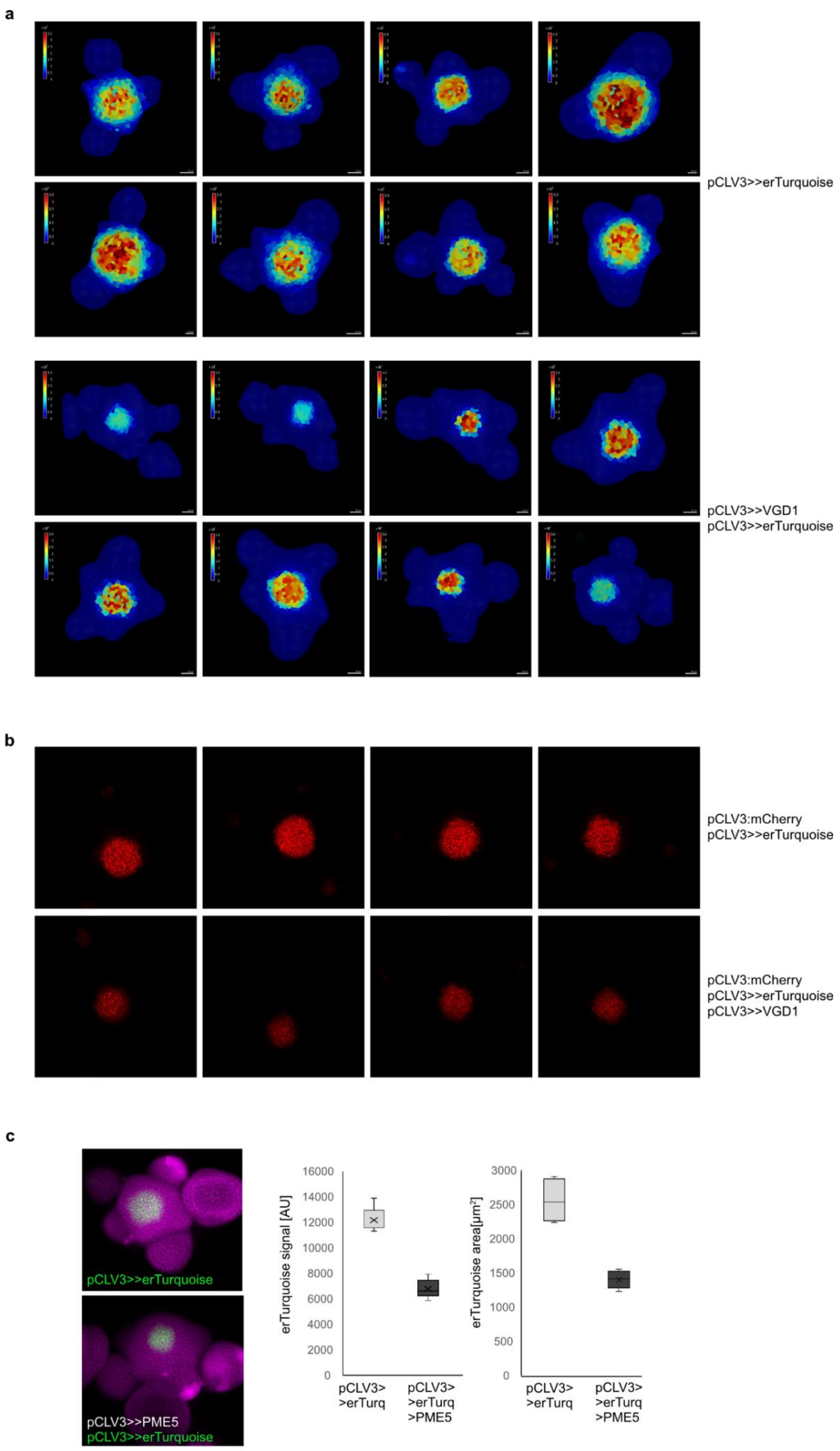
a
pCLV3>>erTurquoise
pCLV3>>VGD1
pCLV3>>erTurquoise
b
pCLV3:mCherry
pCLV3>>erTurquoise
pCLV3:mCherry
pCLV3>>erTurquoise
pCLV3>>VGD1
c
pCLV3>>erTurquoise
pCLV3>>PME5
pCLV3>>erTurquoise
erTurquoise signal [AU]
16000
14000
12000
10000
8000
6000
4000
2000
0
pCLV3> >erTurq
pCLV3> >erTurq >PME5
erTurquoise area[μm²]
3000
2500
2000
1500
1000
500
0
pCLV3> >erTurq
pCLV3> >erTurq >PME5

**Supplemental Figure 5.** Effects of PME expression in stem cells. a) Representative meristem projections after MorphographX segmentation and projection of the pCLV3>>erTurquoise signal. Meristems were images 72 hours after DEX induction. b) . a) Representative meristem projections after MorphographX segmentation and projection of the pCLV3:mCherry-NLS signal. Meristems were images 72 hours after DEX induction. c) Effect of pCLV3>>PME5 expression on mTurquoise signal and domain size.

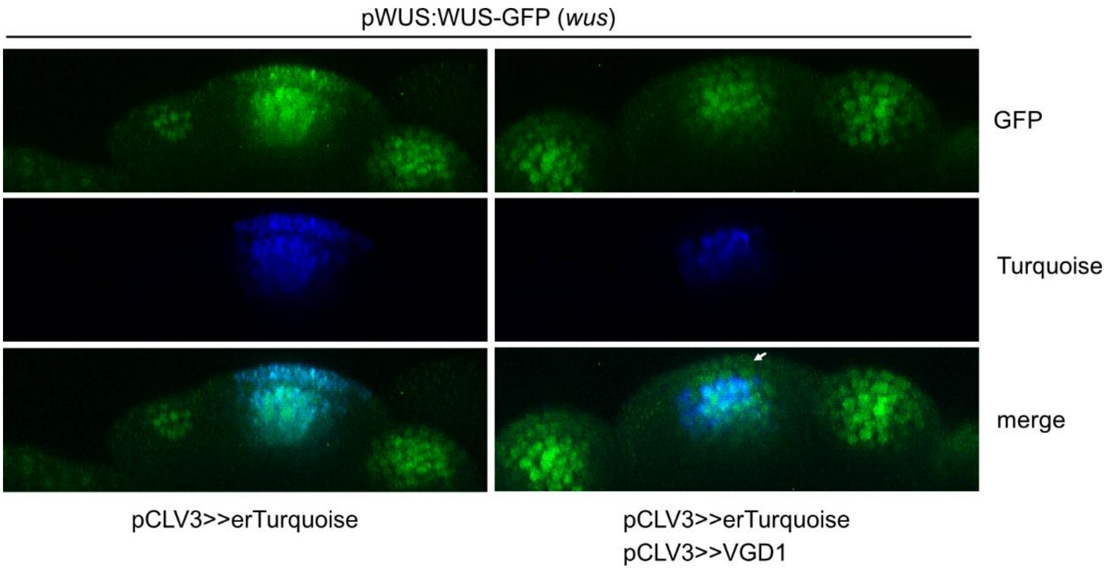


**Supplemental Figure 6.** Cross sections of pWUS:WUS-GFP signal in the indicated bakgrounds. Lines were generated by crossing the WUS rescue line described in (Daum et al., 2014) with the pCLV3>>erTurquoise and the pCLV3>>erTurquoise >>VGD1 lines, respectively. Plants were homozygous for all loci.

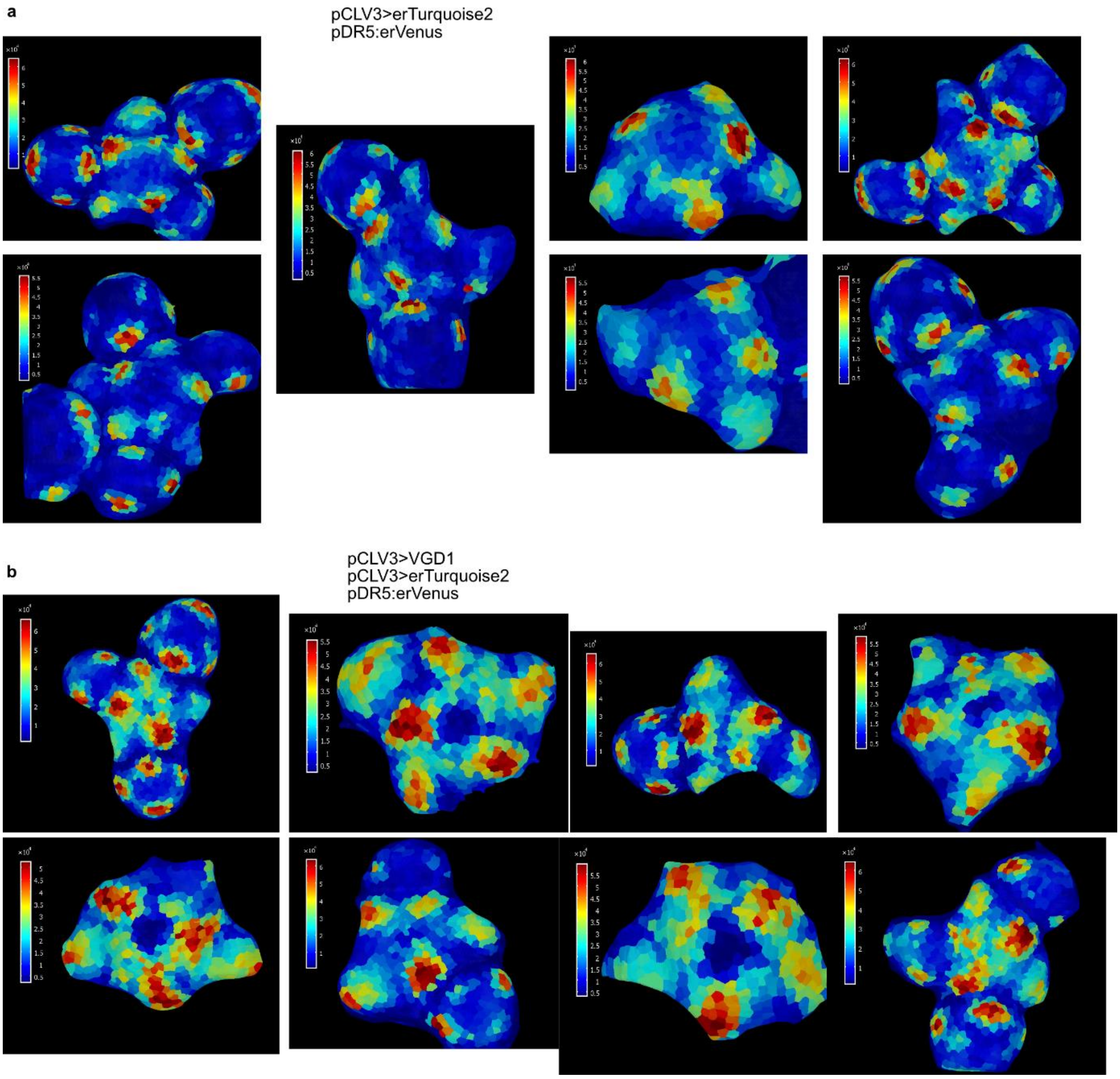


**Supplemental Figure 7.** Effects of PME expression in stem cells. a) Representative meristem projections after MorphographX segmentation and projection of the pDR5:erCitirne signal. Meristems were images 72 hours after DEX induction.
