## Supplemental Table 1 for "The Cell Wall Controls Stem Cell Fate in the Arabidopsis Shoot Apical Meristem"

**Table S1:** Mutants and transgenic lines used in this study

| Mutant/transgenic line | |
|---|---|
| *pCLV3>>mTurqouise* | Schürholz et al., 2018 |
| *pCLV3>>mTurqouise, >>VGD1* | This study |
| *pCLV3>>mTurqouise, >>VGD1; pCLV3:mCherry-NLS* | This study |
| *pCLV3>>mTurqouise, >>VGD1; pSVP:3xGFP-NLS* | This study |
| *pCLV3>>mTurqouise, >>PME5* | This study |
| *pWUS:WUS-GFP (wus)* | Daum et al., 2014 |
| pPME5:erCitrine | This study |
| pPME5:erCitrine_WUSm | This study |
| pPME5:erCitrine $\Delta$ | This study |
| *pPMEPCRF:erCitrine* | This study |
| pUB10:mCherry-GR-WUS | Ma et al., 2019 |
| P35S:VGD1 | Wolf and Greiner 2012 |
| *pDR5:erYFP* | Ma et al., 2019 |

**Gadeyne, A., Sanchez-Rodriguez, C., Vanneste, S., Di Rubbo, S., Zauber, H., Vanneste, K., Van Leene, J., De Winne, N., Eeckhout, D., Persiau, G., Van De Slijke, E., Cannoot, B., Vercruysse, L., Mayers, J.R., Adamowski, M., Kania, U., Ehrlich, M., Schweighofer, A., Ketelaar, T., Maere, S., Bednarek, S.Y., Friml, J., Gevaert, K., Witters, E., Russinova, E., Persson, S., De Jaeger, G. and Van Damme, D.** (2014) The TPLATE adaptor complex drives clathrin-mediated endocytosis in plants. *Cell*, **156**, 691-704.

**Holzwart, E., Huerta, A.I., Glockner, N., Garnelo Gomez, B., Wanke, F., Augustin, S., Askani, J.C., Schurholz, A.K., Harter, K. and Wolf, S.** (2018) BRI1 controls vascular cell fate in the Arabidopsis root through RLP44 and phytosulfokine signaling. *Proc Natl Acad Sci U S A*.

**Jaillais, Y., Belkhadir, Y., Balsemao-Pires, E., Dangl, J.L. and Chory, J.** (2011) Extracellular leucine-rich repeats as a platform for receptor/coreceptor complex formation. *Proc Natl Acad Sci U S A*, **108**, 8503-8507.

**Kutschmar, A., Rzewuski, G., Stuhrwohldt, N., Beemster, G.T., Inze, D. and Sauter, M.** (2009) PSK-alpha promotes root growth in Arabidopsis. *New Phytol*, **181**, 820-831.

**Wolf, S., van der Does, D., Ladwig, F., Sticht, C., Kolbeck, A., Schurholz, A.K., Augustin, S., Keinath, N., Rausch, T., Greiner, S., Schumacher, K., Harter, K., Zipfel, C. and Hofte, H.** (2014) A receptor-like protein mediates the response to pectin modification by activating brassinosteroid signaling. *Proc Natl Acad Sci U S A*, **111**, 15261-15266.
