## Supplemental Table 2 for "The Cell Wall Controls Stem Cell Fate in the Arabidopsis Shoot Apical Meristem"

**Table S2.** Oligonucleotides used in this study. Oligos used for GreenGate Cloning (Lampropoulos et al., 2013 are listed in Supplemental Table 3.

| *Primer No.* | *Primer name* | *Sequence (5′ → 3′)* | *target* |
|---|---|---|---|
| SW2711 | KO_p[PME5]_Wuschel_Fw | ATATATGGTCTCcAGGTTGCTTAAGAAGCAGTGTGTCCG | CRISPR deletion of PME5 WUS BM |
| SW2712 | KO_p[PME5]_Wuschel_rv | ATTATTGGTCTCaACCTCCTTTTGGGGTCCTACTTCC | CRISPR deletion of PME5 WUS BM |
| SW3224 | PMEPCRF_ChIP_qPCR_1F | TCGACCGTTGGATCGATAC | ChIP-PCR |
| SW3225 | PMEPCRF_ChIP_qPCR_1R | GTTGGTGGGCGATGTAGAT | ChIP-PCR |
| SW3226 | PMEPCRF_ChIP_qPCR_2F | AGAACAGGAAGAGTCAAGGC | ChIP-PCR |
| SW3227 | PMEPCRF_ChIP_qPCR_2R | CGTATGGTAATGGAGGACT | ChIP-PCR |
| SW3228 | PMEPCRF_ChIP_qPCR_3F | AAAGTCACGCGGAGAACTAG | ChIP-PCR |
| SW3229 | PMEPCRF_ChIP_qPCR_3R | GACTTATGGTAGAGGACAGCT | ChIP-PCR |
| SW3304 | PME3_ChIP_1F | cacgagtggatgagccataa | ChIP-PCR |
| SW3305 | PME3_ChIP_1R | ctgagtcatgtgagtgtctc | ChIP-PCR |
| SW3306 | PME3_ChIP_2F | atccacataacgtgactcgg | ChIP-PCR |
| SW3307 | PME3_ChIP_2R | cgcacatgtagcctagtagt | ChIP-PCR |
| SW3308 | PME34_ChIP_1F | aggctgtctatagatttcgg | ChIP-PCR |
| SW3309 | PME34_ChIP_1R | gctttctttagtggtccat | ChIP-PCR |
| SW3310 | PME34_ChIP_2F | agtgtccaatagattctcc | ChIP-PCR |
| SW3311 | PME34_ChIP_2R | ggttcgagtacatactact | ChIP-PCR |
| SW3312 | PME44_ChIP_1F | gtaccagcgaaacccacatta | ChIP-PCR |
| SW3313 | PME44_ChIP_1R | tggctcacgtgacctagtta | ChIP-PCR |
| SW3314 | PME44_ChIP_2F | tctctttatggcccaatggc | ChIP-PCR |
| SW3315 | PME44_ChIP_2R | cacaccggaagcagctgttt | ChIP-PCR |
| SW3304 | PME3_ChIP_1F | cacgagtggatgagccataa | ChIP-PCR |
| SW3305 | PME3_ChIP_1R | ctgagtcatgtgagtgtctc | ChIP-PCR |
| SW3306 | PME3_ChIP_2F | atccacataacgtgactcgg | ChIP-PCR |

| | | | |
|---|---|---|---|
| SW3307 | PME3_ChIP_2R | cgcacatgtagcctagtagt | ChIP-PCR |
