## Supplemental Table 3 for "The Cell Wall Controls Stem Cell Fate in the Arabidopsis Shoot Apical Meristem"

| | | | |
|---|---|---|---|
| **pSW930** | pOp6:VGD1 | | |
| pSW180 | pOp6 (pGGA016) | Lampropoulos et al., 2013 | |
| pSW922 | VGD1_BD+STOP | aacaGGTCTCaAACAATGATTGGAAAAGTTGTGGTCTC | aacaGGTCTCtGCAGTTATAATCCAAGCGTGACGGG |
| pSW186 | UBQ10 terminator (pGGE009) | Lampropoulos et al., 2013 | |
| pSW319 | pMAS:BastaR:tMAS ( pGGF001) | Lampropoulos et al., 2013 | |
| pGGZ001 | Destination vector | Lampropoulos et al., 2013 | |
| **pSW605** | pOp6:PME5 | | |
| pSW180 | pOp6 (pGGA016) | Lampropoulos et al., 2013 | |
| pSW182 | B-Dummy (pGGA022) | Lampropoulos et al., 2013 | |
| pSW445 | PME5 | AACAGGTCTCAGGCTATGGCGCAACTTACTAATTCC | AACAGGTCTCACTGATTAAGCATCTCGAGGAGCGA |
| pSW184 | D-dummy (pGGD002) | Lampropoulos et al., 2013 | |
| pSW186 | UBQ10 terminator (pGGE009) | Lampropoulos et al., 2013 | |
| pSW319 | pMAS:BastaR:tMAS ( pGGF001) | Lampropoulos et al., 2013 | |
| pGGZ001 | Destination vector | Lampropoulos et al., 2013 | |
| **pSW1008** | pPME5:erCitrine | | |
| pSW810 | pPME5 | aacaGGTCTCaACCTcatccgcaacgatagattat | aacaGGTCTCtTGTTtggcttgtgagaaaggaaac |
| pSW548 | Signal Peptide (pGGB006) | Lampropoulos et al., 2013 | |
| pSW962 | GAGAGA-mCitrine (pPD159) | Denninger et al., 2019 | |
| pSW550 | HDEL (pGGD008) | Lampropoulos et al., 2013 | |
| pSW965 | PME5 term | AACAGGTCTCACTGCCCAACTTCAAACCTTGGCGG | AACAGGTCTCTTAGTGTGACCAGGAACTGAAAATTTC |
| pSW319 | pMAS:BastaR:tMAS ( pGGF001) | Lampropoulos et al., 2013 | |
| pGGZ001 | Destination vector | Lampropoulos et al., 2013 | |
| **pSW1095** | pPME5:erCitrine_WUSm | | |
| pSW1094 | pPME5_WUSm | CAAAAGGAGGTTaACGTaATGCTTAAG | CTTAAGCATtACGTtAACCTCCTTTTG |
| pSW548 | Signal Peptide (pGGB006) | Lampropoulos et al., 2013 | |
| pSW962 | GAGAGA-mCitrine (pPD159) | Denninger et al., 2019 | |

| | | | |
|---|---|---|---|
| pSW550 | HDEL (pGGD008) | Lampropoulos et al., 2013 | |
| pSW965 | PME5 term | As above | |
| pSW319 | pMAS:BastaR:tMAS ( pGGF001) | Lampropoulos et al., 2013 | |
| pGGZ001 | Destination vector | Lampropoulos et al., 2013 | |
| **pSW1209** | pSVP:3xGFP-NLS | | |
| pSW1024 | pSVP | aacaGGTCTCaACCTgaaagaagcctaaaatggc | aacaGGTCTCtTGTTCACAACGAACAAAAAAACCC |
| pSW182 | B-Dummy (pGGA022) | Lampropoulos et al., 2013 | |
| pSW322 | 3xGFP (pGGC025) | Lampropoulos et al., 2013 | |
| pSW183 | linker-NLS (pGGD007) | Lampropoulos et al., 2013 | |
| pSW186 | UBQ10 terminator (pGGE009) | Lampropoulos et al., 2013 | |
| pSW187 | pNOS:KanR:tNOS (pGGF007) | Lampropoulos et al., 2013 | |
| pGGZ001 | Destination vector | Lampropoulos et al., 2013 | |
| **pSW1009** | pPCRF:erCitrine | | |
| pSW808 | pPMEPCRF | aacaGGTCTCaACCTtaattatatatataacgttctttg | aacaGGTCTCtTGTTagttttttttttttttttggttta |
| pSW548 | Signal Peptide (pGGB006) | Lampropoulos et al., 2013 | |
| pSW962 | GAGAGA-mCitrine (pPD159) | Denninger et al., 2019 | |
| pSW550 | HDEL (pGGD008) | Lampropoulos et al., 2013 | |
| pSW986 | PMEPCRFterm | aacaggtctcttagtatcaaaatcagccaacccttcaaaccta | aacaggtctcactgcattgtctcagccttgtatgttacg |
| pSW319 | pMAS:BastaR:tMAS ( pGGF001) | Lampropoulos et al., 2013 | |
| pGGZ001 | Destination vector | Lampropoulos et al., 2013 | |
| **pYM92** | pOp6: NSlmb-vhhGFP4 | | |
| pSW180 | pOp6 (pGGA016) | Lampropoulos et al., 2013 | |
| pSW182 | B-Dummy (pGGA022) | Lampropoulos et al., 2013 | |
| pYM | NSlmb-vhhGFP4 | Ma et al., 2019 | |
| pSW184 | D-dummy (pGGD002) | Lampropoulos et al., 2013 | |
| pGGE001 | Rbcs terminator | Lampropoulos et al., 2013 | |
| pGGF005 | pUBQ10::HygrR::tOCS | Lampropoulos et al., 2013 | |
| pGGZ001 | Destination vector | Lampropoulos et al., 2013 | |

**Table S2**. Overview of constructs generated with GreenGate cloning (Lampropoulos et al., 2013) and the primers used to generate modules, where appropriate.
